# Interleukin-34 Promotes Activation of Disease-associated Microglia-like Cells and Attenuates Disease Progression in West Nile Virus Infection

**DOI:** 10.64898/2026.08.04.742676

**Authors:** Passawat Thammahakin, Keisuke Maezono, Thuy Thi Ngoc Duong, Haruto Eguchi, Savit Promwattanapan, Michihito Sasaki, Hiroaki Kariwa, Yasuko Orba, Shintaro Kobayashi

## Abstract

West Nile virus (WNV) is a mosquito-borne *orthoflavivirus* that causes severe encephalitis, for which no approved antiviral therapies or human vaccines are currently available. Disease-associated microglia (DAM) represent a microglial state associated with neuroprotective functions in neurodegenerative diseases. We previously showed that DAM-like cells are localized in the vicinity of WNV-infected cells in the mouse brain and surmised that these cells may respond to WNV-infected cells. However, the functional significance of this spatial association has since remained unclear. Given the previous reports linking interleukin-34 (IL-34) to the activation of DAM-like cells, we investigated whether IL-34 promotes DAM-like responses and whether these responses are associated with protection against WNV infection. Transcriptomic analysis of IL-34–treated HMC3 human microglial cells revealed upregulation of markers characteristic of DAM, including *TREM2*, *APOE*, *FABP5*, and *FTH1*. IL-34–treated HMC3 cells suppressed WNV replication in co-culture with WNV-infected SH-SY5Y human neuroblastoma cells apparently in a cell–cell contact-dependent manner, independent of secreted factors. In WNV-infected mice, IL-34 administration improved survival, reduced viral titers, and decreased neuronal apoptosis. IL-34 also increased the abundance of CD11c- and SPP1-positive DAM-like cells, predominantly in the vicinity of WNV-infected cells, in the mouse brain. These findings indicate that IL-34 promotes the activation of DAM-like cells and enhances protective responses in WNV encephalitis.

**IMPORTANCE:** West Nile virus (WNV) is a major cause of viral encephalitis worldwide, with no approved antiviral treatment or human vaccine currently available. Microglia, the resident immune cells of the brain, play key roles in responding to viral infection. However, the mechanism through which specific microglial activation states contribute to protection in viral encephalitis remains poorly understood. In this study, we show that interleukin-34 promotes microglial responses resembling disease-associated microglia (DAM) and is associated with enhanced protection against WNV infection in cell culture and mouse models. The study findings suggest that activation of DAM-like cells may contribute to protective host responses against neurotropic viral infection and provide new insights into the role of specific microglial states in viral encephalitis.

## INTRODUCTION

West Nile virus (WNV) is a mosquito-borne *orthoflavivirus*, which is transmitted to humans through the bite of infected Culex mosquitoes. Upon infection, WNV enters the bloodstream and invades the central nervous system (CNS) by disrupting the blood–brain barrier (BBB) and trafficking within infected immune cells (1, 2). Since its emergence and subsequent spread in North America in 1999, WNV has become the most prevalent mosquito-borne virus causing neurological disease worldwide (3). Although most infections are asymptomatic, approximately 1% of infected individuals develop severe neurological complications, such as meningitis, encephalitis, and acute flaccid paralysis (3–5).Within the CNS, WNV targets neurons and induces apoptosis (6, 7), while triggering glial activation and neuroinflammation that further exacerbate the CNS pathology (8). No approved antiviral therapy or vaccine exists for WNV infection in humans (9, 10), warranting the need for a better understanding of WNV pathogenesis to inform future therapeutic strategies. As resident immune cells of the CNS, microglia play essential roles in maintaining brain homeostasis and in responding to pathological stimuli. Following WNV entry into the CNS, microglia become activated and exhibit diverse functional phenotypes, which can contribute to neuroprotective or neurotoxic responses (8, 11). Among these activation states, a transcriptomically unique microglial phenotype, represented by disease-associated microglia (DAM), characterized by reduced expression of homeostatic genes and increased expression of genes involved in phagocytosis and immune activation, including *TREM2*, *APOE*, *SPP1*, and *CD11c*, has been identified (12). Although DAM play protective roles in neurodegenerative diseases, such as Alzheimer’s disease (AD) and amyotrophic lateral sclerosis (ALS) (13, 14), their function in neurotropic viral infections remains unclear. We previously reported the presence of DAM-like cells in WNV-infected mice (15) and suggested that DAM may be induced as part of the microglial response to WNV infection. However, it has remained unclear whether DAM contribute to protection or pathology during WNV infection, or whether promoting their induction attenuates WNV-induced neuropathology.

Colony-stimulating factor 1 receptor (CSF1R) is a key regulator of microglial proliferation, migration, differentiation, and survival (16). CSF1R signaling is activated by two distinct ligands—CSF-1 and interleukin-34 (IL-34)—both of which contribute to the maintenance of microglial homeostasis in the CNS (17, 18). Although both ligands signal through CSF1R, they differ in their expression patterns and effects on microglial function. CSF-1 is ubiquitously expressed throughout the body, whereas IL-34 expression in the CNS predominantly occurs in neurons, which is suggestive of a more specialized role for IL-34 in regulating microglial activity within the brain (17–19). Notably, IL-34–deficient mice exhibit reduced microglial numbers and impaired CNS defenses against viral infection, which highlights the importance of IL-34–mediated signaling in microglia-dependent neuroprotection (17, 20). IL-34 is also linked to TREM2-mediated microglial responses (21, 22). TREM2 is a key regulator of DAM transition (12, 23). These observations raise the possibility that IL-34 may influence DAM-like cell responses during viral infection.

In this study, we aimed to investigate whether IL-34 promotes the activation of DAM-like cells and whether this response is associated with protection in WNV infection. We conducted transcriptomic analysis of IL-34–treated human microglial cells to characterize the expression of genes characteristic of the DAM phenotype, evaluated the antiviral effects of IL-34–treated microglia in co-culture with WNV-infected cells, and examined the effect of IL-34 administration in a murine model of WNV encephalitis. Our findings show that IL-34 promotes the activation of DAM-like cells and enhances protective responses against WNV infection, providing novel insights into how microglial activation states contribute to host defense against neurotropic viral infection.

## RESULTS

### IL-34 induces a DAM-like transcriptional signature in human microglial cell line

IL-34 treatment of microglia has been reported to activate neuroprotective microglial responses through TREM2 signaling, which is a key pathway of DAM induction (12, 21, 24). To examine whether IL-34 treatment induces a DAM-like state in HMC3, a human microglial cell line, the cells were treated with recombinant IL-34 (200 ng/mL) for 24 h and processed for bulk RNA sequencing. Principal component analysis (PCA) revealed clear separation between IL-34– and vehicle-treated cells, confirming distinct transcriptional profiles of the two groups (Fig. 1A). Differential gene expression analysis revealed 473 upregulated and 293 downregulated genes in IL-34–treated cells compared with vehicle control–treated cells (Fig. 1B). Several markers related to the DAM phenotype, including *TREM2*, *APOE*, *FABP5*, and *FTH1*, showed a trend toward upregulation (Fig. 1C). Furthermore, gene ontology (GO) enrichment analysis of upregulated genes revealed significant enrichment of chemotaxis-related biological processes, including regulation of chemotaxis and cell chemotaxis, driven by upregulation of *TMSB4X*, *MIF*, *HSPB1*, *GSTP1*, *CREB3*, and *DDT* (Fig. 1B and S1A). In addition, GO enrichment analysis of downregulated genes was performed and the top 10 differentially expressed genes (DEGs) were identified (Fig. S1B and S1C). To validate whether these transcriptomic findings indicated a DAM-like state, reverse transcription-quantitative PCR (RT-qPCR) was performed on selected markers characteristic of DAM. IL-34–treated HMC3 cells showed significantly increased mRNA expression of *TREM2*, *SPP1*, and *CD11c* compared with vehicle-treated controls (Fig. 1D). Moreover, immunofluorescence staining confirmed an increase in the number of CD11c-positive cells in IL-34–treated HMC3 cells compared with that in vehicle-treated controls (Fig. 1E–F). These findings indicate that IL-34 induces a DAM-like transcriptional pattern in human microglial cell line, characterized by upregulation of DAM markers.

**FIG 1.**
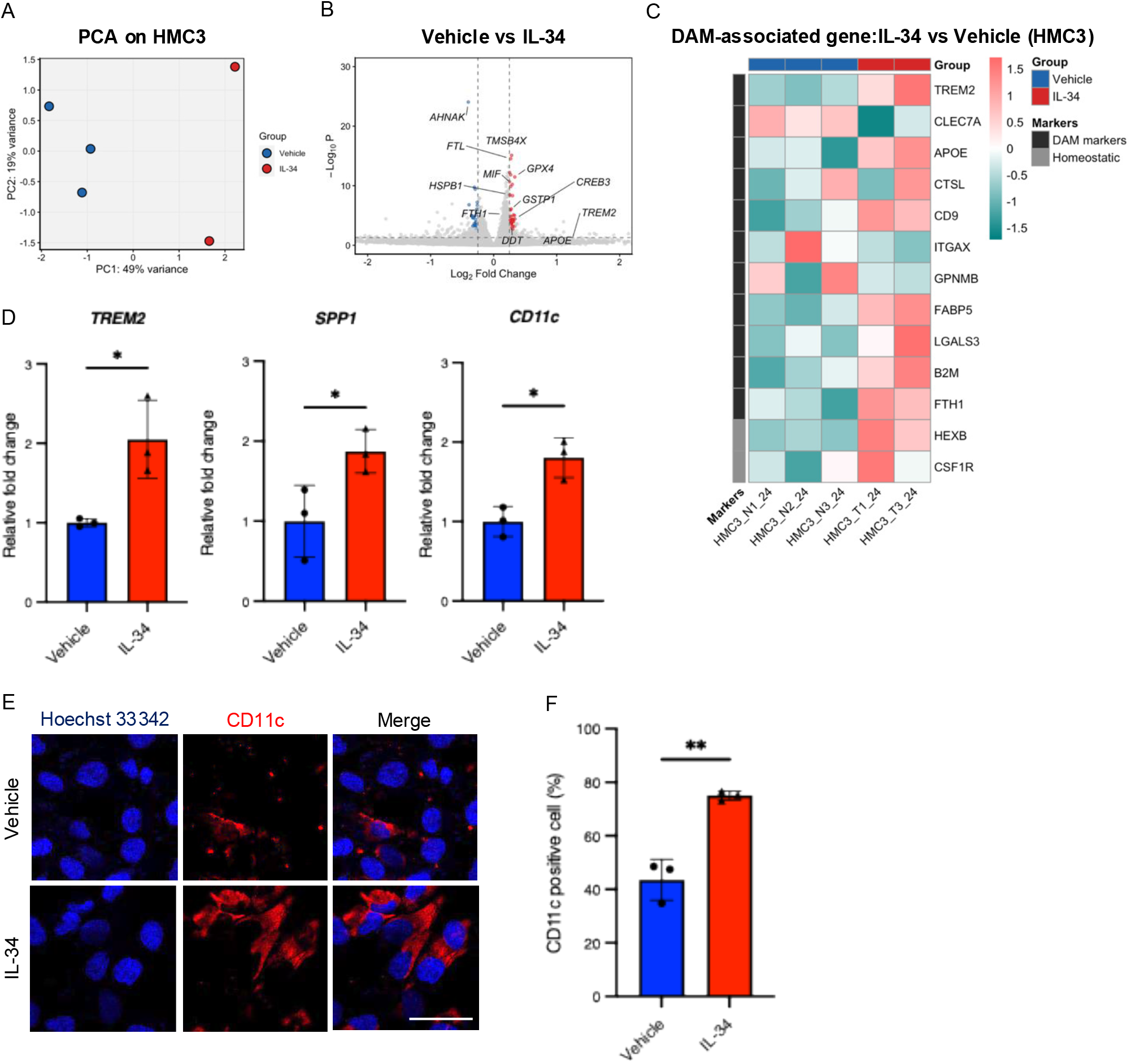
Transcriptomic analysis of the expression of genes associated with disease-associated microglia (DAM) in interleukin (IL)-34–treated HMC3 cells. (A) Principal component analysis (PCA) of transcriptomic profiles of IL-34–treated (*n* = 2) and vehicle-treated (*n* = 3) HMC3 cells. Volcano plot of differentially expressed genes (DEGs) in IL-34–treated versus vehicle-treated HMC3 cells. Red dots indicated upregulated genes (log2 fold change >0.25, FDR-adjusted *P*-value <0.05) and blue dots indicated downregulated genes (log2 fold change <−0.25, FDR-adjusted *P*-value <0.05). Selected genes are labeled. (C) Heatmap of the expression of DAM-associated and homeostatic marker genes across IL-34– and vehicle-treated HMC3 cells. Color scale represents row-normalized expression values. (D) Relative mRNA expression of *TREM2*, *SPP1*, and *CD11c* in IL-34–treated to vehicle-treated HMC3 cells assessed via RT-qPCR. Data represent the mean ± SD (*n* = 3 per group). (E) Representative immunofluorescence images of CD11c in vehicle- or IL-34–treated HMC3 cells. Scale bar, 50 μm. (F) Quantification of CD11c-positive cells as a percentage (manually counting >100 cells per sample). Data represent the mean ± SD (*n* = 3 per group). Statistical significance was determined by Student’s *t*-test. *, *P* < 0.05; **, *P* < 0.01.

### IL-34–treated HMC3 cells restrict WNV replication in neuronal SH-SY5Y cells

The close spatial association between DAM and WNV-infected cells has been reported, and their interaction may contribute to WNV pathogenesis (15). As IL-34 induced a DAM-like transcriptional pattern in HMC3 cells, we investigated whether IL-34–treated microglia affected WNV replication in neuronal cells. SH-SY5Y cells were highly susceptible to WNV infection, with widespread detection of viral antigen, whereas the infection in HMC3 cells was limited to only a few cells (Fig. S2). First, to examine whether DAM-like cells contribute to reduced viral replication in neuronal cells, IL-34–treated HMC3 cells were co-cultured with WNV-infected SH-SY5Y cells (co-culture) (Fig. 2A). Co-culture of IL-34–treated HMC3 cells with WNV-infected SH-SY5Y cells significantly reduced viral titers at 12 h post-infection (hpi), with a greater reduction observed at 24 hpi compared with that in vehicle-treated HMC3 cells (Fig. 2A). Second, to determine whether soluble factors derived from IL-34–treated HMCs are involved in suppressing WNV replication, supernatant from IL-34–treated HMC3 cells was transferred to WNV-infected SH-SY5Y cells (supernatants) (Fig. 2B). The supernatant collected from IL-34–treated HMC3 cells did not affect viral titers at either timepoint (Fig. 2B). Third, to determine whether IL-34 directly affects WNV replication in neuronal cells, SH-SY5Y cells were treated directly with recombinant IL-34 (Fig. 2C). IL-34 treatment of SH-SY5Y cells had no effect on the titer of progeny virus at 12 or 24 hpi (Fig. 2C). To further confirm the close spatial association between DAM-like cells and WNV-infected cells in the co-culture system, their localization was examined via immunofluorescence staining for SPP1 and viral antigen (Fig. 2D). SPP1-positive cells were observed near viral antigen–positive cells. Overall, these findings indicated that IL-34–treated microglia plausibly suppress WNV replication through direct cell–cell contact with infected neuronal cells, rather than through secreted factors.

**FIG 2.**
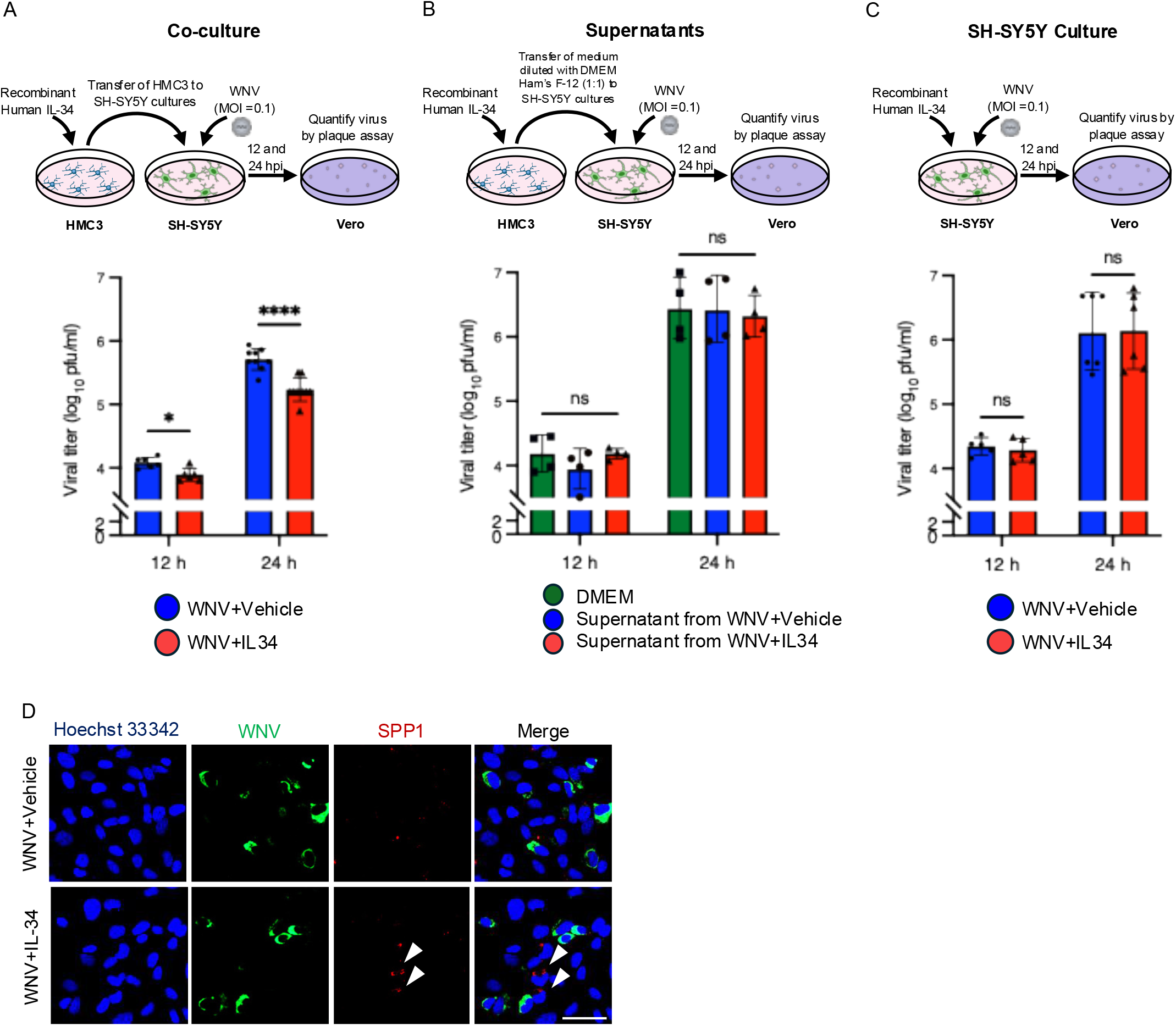
Co-culture with interleukin (IL)-34–treated HMC3 cells restricts West Nile virus (WNV) replication in SH-SY5Y cells. (A) Schematic diagram and viral titers in WNV-infected SH-SY5Y cells at a multiplicity of infection (MOI) of 0.1, cocultured with IL-34– or vehicle-treated HMC3 cells at 12 and 24 h post-infection (hpi). (B) Schematic diagram and viral titers in WNV-infected SH-SY5Y cells at an MOI of 0.1, treated with conditioned medium collected from IL-34– or vehicle-treated HMC3 cells at 12 and 24 hpi. (C) Schematic diagram and viral titers in WNV-infected SH-SY5Y monoculture at an MOI of 0.1, treated with IL-34 (200 ng/mL) or vehicle at 12 and 24 hpi. Data are presented as the mean ± SD (*n* ≥ 4 per group) for panels A to C. (D) Representative immunofluorescence images of WNV (green) and SPP1 (red) staining in co-cultured WNV-infected SH-SY5Y cells with vehicle-treated (WNV+Vehicle) or IL-34–treated (WNV+IL-34) HMC3 cells. White arrowheads indicate SPP1-positive cells. Scale bar, 50 μm. ns, not significant. Statistical significance was determined via one-way ANOVA with Tukey’s post-hoc test. *, *P* < 0.05; ****, *P* < 0.0001.

### IL-34 induces a DAM-like microglial state in the mouse brain

To extend our *in vitro* findings to *in vivo* settings, we examined the effects of IL-34 on microglial responses in the mouse brain. Treatment with recombinant IL-34 induced microglial activation, as evidenced by increased TMEM119-positive cell area (Fig. 3A and 3B). A similar increase was also observed in the brain of mice treated with recombinant CSF-1, which shares the same receptor, CSF1R, with IL-34 (Fig. S3A and S3B) (17). Next, we examined the expression of two DAM markers, CD11c and SPP1, in IL-34–treated mice. The number of CD11c-positive cells was significantly increased following IL-34 treatment compared with that after vehicle treatment (Fig. 3C and 3D). Consistent with these results, *CD11c* mRNA expression in TMEM119-sorted cells was upregulated in IL-34–treated mice compared with that in vehicle-treated controls (Fig. 3E). As observed for CD11c, the number of SPP1-positive cells was increased following IL-34 treatment (Fig. 3F and 3G). In contrast, the number of CD11c- or SPP1-positive cells did not increase following recombinant CSF-1 treatment (Fig. S3C and S3D). These findings indicated that IL-34 administration promotes the expansion of DAM-like cells in the mouse brain.

**FIG 3.**
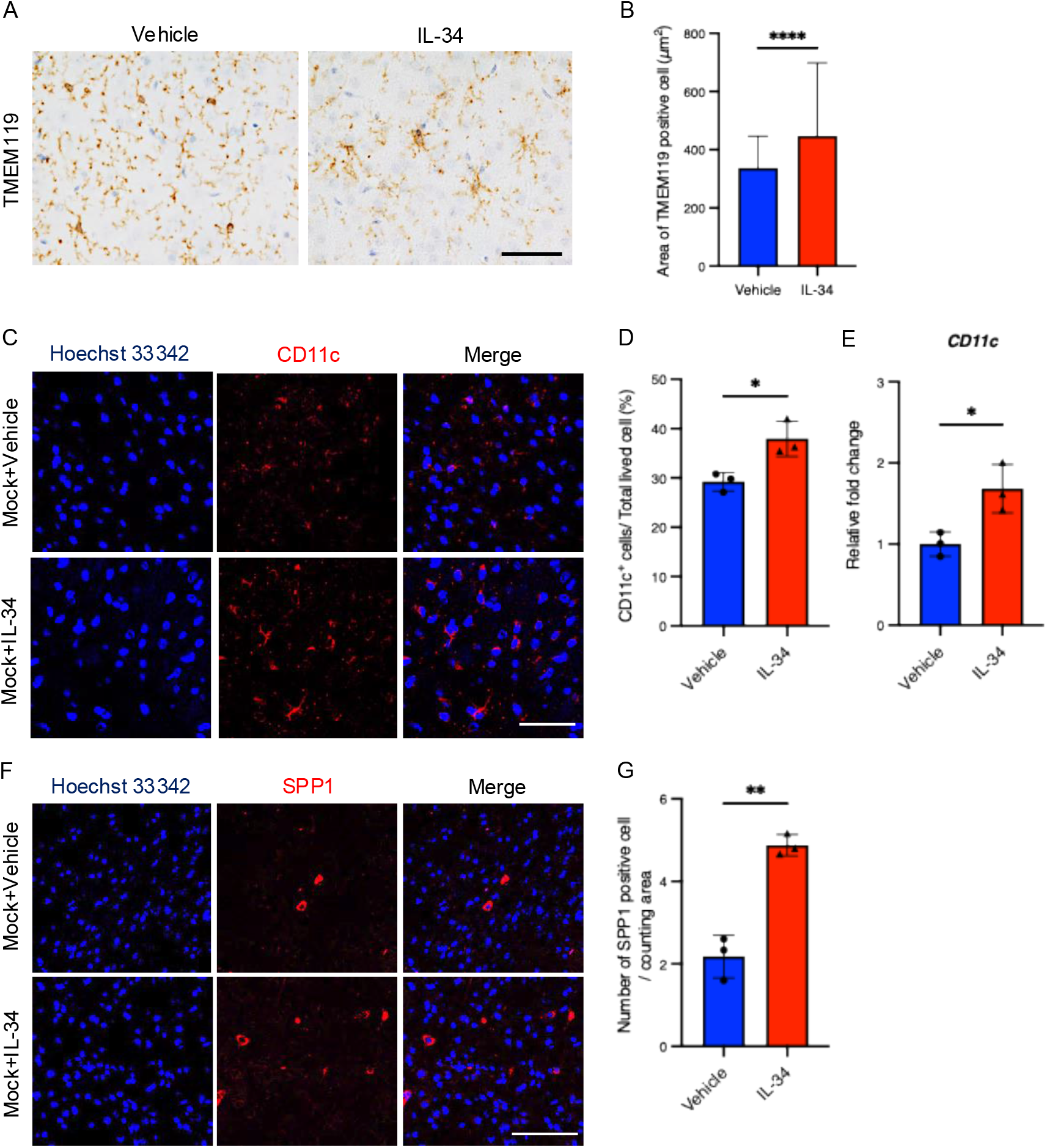
Analysis of interleukin (IL)-34–induced disease-associated microglia (DAM)-like microglia in vivo. (A) Representative immunohistochemistry images of TMEM119 expression in brain sections from mice treated with vehicle or IL-34. Scale bar, 50 μm. (B) Quantification of the average area of TMEM119-positive cells representing microglial size. Data represent the mean ± SD from three fields per mouse (*n* = 3 mice per group). (C) Representative immunofluorescence images of brain sections from vehicle- and IL-34–treated mice stained for CD11c. Scale bar, 50 μm. (D) Quantification of CD11c-positive cells as a percentage of total live cells. Data represent the mean ± SD (*n* = 3 mice per group). (E) Relative mRNA expression of *CD11c* in TMEM119-sorted cells from vehicle- and IL-34–treated uninfected mice assessed via RT-qPCR. Data represent the mean ± SD (*n* = 3 mice per group). (F) Representative immunofluorescence images of brain sections from vehicle- and IL-34–treated mice stained for SPP1. Scale bar, 100 μm. (G) Quantification of SPP1-positive cells per counting area. Data are presented as the mean ± SD from five fields per mouse (*n* = 3 per group). Statistical significance was determined via Student’s *t*-test. *, *P* < 0.05; **, *P* < 0.01; ****, *P* < 0.0001.

### IL-34 treatment mitigates WNV disease progression in WNV-infected mice

Based on our findings that IL-34 induces a DAM-like state and reduces viral replication *in vitro*, we next investigated whether IL-34 treatment protects against WNV disease progression *in vivo*. Mice were intracranially inoculated with 10 plaque-forming units (pfu) of WNV and subsequently treated with 100 ng of recombinant IL-34, CSF-1, or vehicle on the following day (Fig. 4A). IL-34–treated WNV-infected mice showed a significantly higher survival rate than vehicle-treated mice, whereas no significant difference in survival was observed between CSF-1– and vehicle-treated mice (Fig. 4B). Additionally, loss in body weight at 7 dpi was attenuated in IL-34–treated mice compared with that in vehicle-treated mice (Fig. 4C).

**FIG 4.**
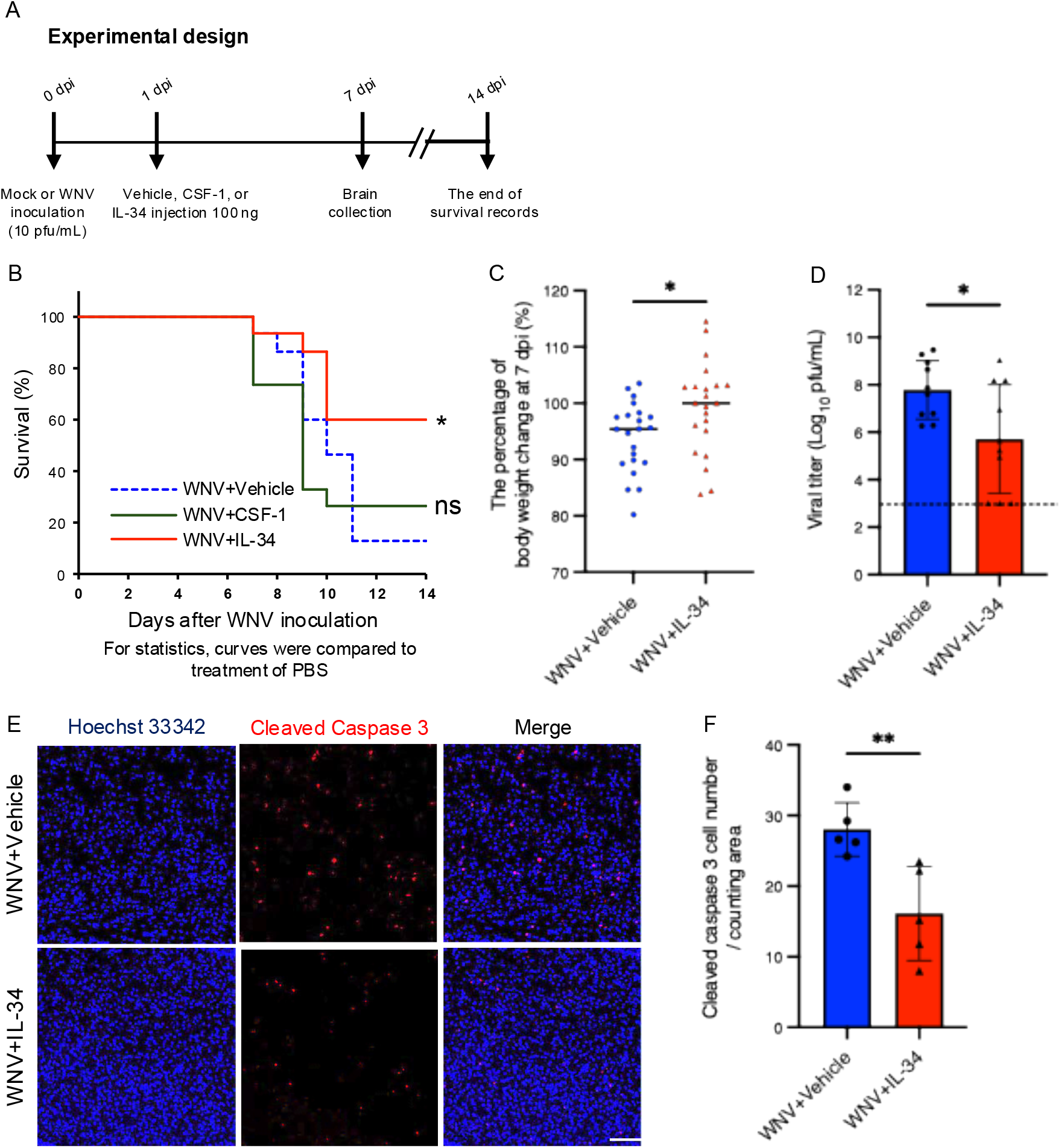
Survival and pathological analyses of protective effects of interleukin (IL-34) against West Nile virus (WNV) disease progression. (A) Schematic of the experimental design. Mice were intracranially inoculated with WNV (10 pfu/mL) on day 0 and administered vehicle, CSF-1, or IL-34 (100 ng in 10 µL PBS) on day 1. Brain tissues were collected from a group of mice at 7 days post-infection (dpi) and survival of another subset of mice was recorded until 14 dpi. (B) Kaplan–Meier survival curves for WNV-infected mice treated with vehicle, CSF-1, or IL-34. Survival curves were compared with those for the vehicle-treated group. (C) Percentage change in body weight on 7 dpi relative to initial body weight of mice treated with vehicle or IL-34 (*n* ≥ 19 per group). (D) WNV titers in brain tissues collected on 7 dpi, determined via plaque assay on Vero cells. Data are shown as log10 (pfu/mL) of viral titers from homogenized brain tissue (*n* = 8 per group). The dotted line indicates the limit of detection. (E) Representative immunofluorescence images of cleaved caspase-3 (red) in brain sections from WNV-infected mice treated with vehicle (WNV+Vehicle) or IL-34 (WNV+IL-34). Scale bar, 100 μm. (F) Quantification of cleaved caspase-3-positive cells per counting area. Data are presented as the mean ± SD from five fields per mouse (*n* = 5 per group). Statistical significance was determined via the log-rank test for panel B, and Student’s *t*-test for panels C, D, and F. ns, not significant; *, *P* < 0.05; **, *P* < 0.01.

To determine whether the increased survival of IL-34–treated mice was associated with reduced viral replication, WNV titers in the brain tissue at 7 dpi were quantified via plaque assay. Viral titers in IL-34–treated mice were significantly lower than those in vehicle-treated mice, and viral titers in three IL-34–treated mice were below the limit of detection of the plaque assay (Fig. 4D). As neuronal apoptosis is a hallmark of WNV neuropathogenesis (7, 25), the effect of IL-34 treatment on neuronal apoptosis was examined by quantifying cleaved caspase-3–positive cells. The number of cleaved caspase-3–positive cells was significantly reduced in IL-34–treated mice compared with that in vehicle-treated mice (Fig. 4E-F). In contrast, no significant differences in body weight change, viral titer, or number of apoptotic cells were observed in CSF-1–treated mice compared with that in vehicle-treated mice (Fig. S4A-D). These results indicated that IL-34 treatment suppressed disease progression in WNV-infected mice by reducing viral replication and neuronal apoptosis.

### IL-34 treatment expands DAM-like cell populations in the brain of WNV-infected mice

Given that IL-34 promotes the activation of DAM-like cells in uninfected mice (Fig. 3) and improves survival in WNV-infected mice (Fig. 4), we next investigated whether DAM-like cells contribute to IL-34–mediated protection by assessing their expansion in the brain of WNV-infected mice treated with IL-34. To characterize and visualize DAM-like cells, brain sections from WNV-infected mice were stained for TMEM119, CD11c, and viral antigen. CD11c-positive and TMEM119-weakly positive cells, consistent with a DAM-like phenotype (12), were observed in both vehicle- and IL-34–treated groups (Fig. 5A). The number of these DAM-like cells was significantly increased in IL-34–treated WNV-infected mice compared with that in vehicle-treated mice (Fig. 5B and 5C). We further examined the spatial association between DAM-like cells and WNV antigen–positive cells. In the brain of WNV-infected mice treated with IL-34, the numbers of CD11c- and SPP1-positive DAM-like cells were significantly increased (Fig. 5D, 5E, 5G, and 5H). In addition, these cells were more frequently observed to be in close proximity of WNV antigen–positive cells in IL-34–treated mice (Fig. 5D, 5F, 5G, and 5I). In contrast, CSF-1 treatment did not increase the number of CD11c-positive DAM-like cells and the frequency of their localization in close proximity of WNV antigen–positive cells (Fig. S5A–C). Taken together, these findings indicated that IL-34 promotes the expansion of DAM-like cells and their close spatial association with WNV-infected cells in WNV-infected mouse brain, which may contribute to IL-34–mediated suppression of disease progression.

**FIG 5.**
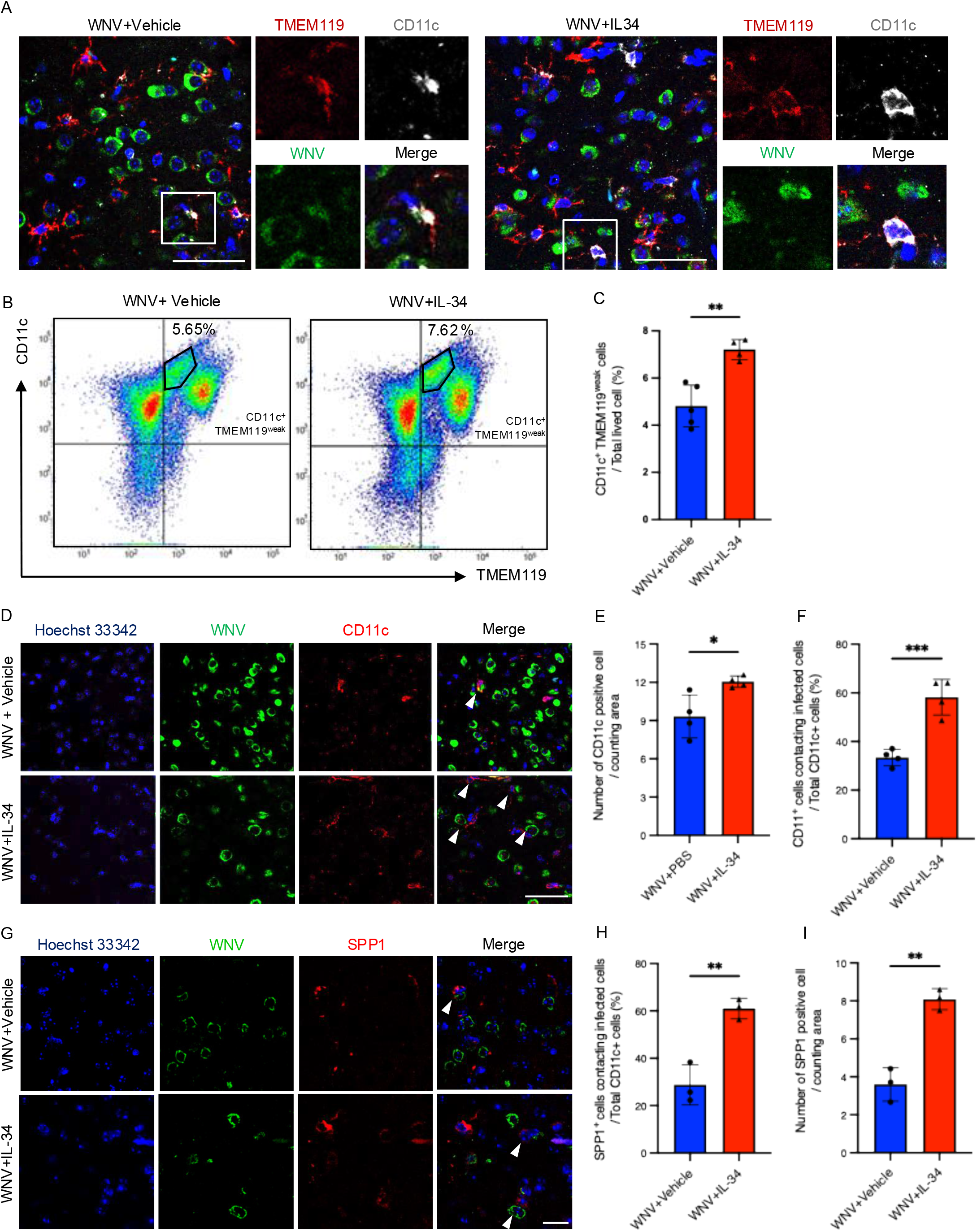
Association between interleukin (IL-34) treatment and DAM-like microglial activation in West Nile virus (WNV)-infected mice. (A) Representative triple immunofluorescence images of TMEM119 (red), CD11c (white), and WNV antigen (green) staining in brain sections from WNV-infected mice treated with vehicle (WNV+Vehicle) or IL-34 (WNV+IL-34). Insets show enlarged views of the areas indicated by white boxes. Scale bar, 50 μm. (B) Representative flow cytometry dot plots of CD11c-positive TMEM119 weak-positive cell populations in WNV-infected mice treated with vehicle or IL-34 (*n* = 4 per group). (C) Quantification of CD11c-positive TMEM119 weak-positive cells as a percentage of total microglia from WNV+Vehicle and WNV+IL-34 groups. Data are presented as the mean ± SD (*n* = 4 per group). (D) Representative immunofluorescence images of CD11c (red) and WNV antigen (green) in brain sections from WNV-infected mice treated with vehicle (WNV+Vehicle) or IL-34 (WNV+IL-34). White arrowheads indicate CD11c-positive cells in contact with WNV-infected cells. Scale bar, 100 μm. (E and F) Quantification of CD11c-positive cells per counting area and percentage of CD11c-positive cells in contact with WNV-infected cells out of total CD11c-positive cells. Data are presented as the mean ± SD from five fields per mouse (*n* = 3 per group). (G) Representative immunofluorescence images of SPP1 (red) and WNV antigen (green) in brain sections from WNV-infected mice treated with vehicle (WNV+Vehicle) or IL-34 (WNV+IL-34). White arrowheads indicate SPP1-positive cells in contact with WNV-infected cells. Scale bar, 50 μm. (H and I) Quantification of SPP1-positive cells per counting area and percentage of SPP1-positive cells in contact with WNV-infected cells out of total SPP1-positive cells. Data are presented as the mean ± SD from five fields per mouse (*n* = 3 per group). Statistical significance was determined via Student’s *t*-test. ns, not significant; *, *P* < 0.05; **, *P* < 0.01; ***, *P* < 0.001.

## DISCUSSION

In this study, we observed that IL-34 treatment promoted the expression of genes characteristic of the DAM phenotype in a human microglial cell line and suppressed WNV replication through attachment with infected neuronal cells. Moreover, IL-34 administration improved survival in WNV-infected mice, accompanied by reduction in viral titers and neuronal apoptosis. IL-34 administration promoted the expansion of DAM-like cells in WNV-infected mouse brain. These findings suggest that IL-34 promotes the induction of a neuroprotective DAM-like phenotype in WNV infection.

Although DAM-like cells have been suggested to play protective roles in neurodegenerative diseases, such as AD and ALS, their function during viral encephalitis, particularly WNV infection, remains poorly understood (12–14). The present study indicates that the induction of DAM-like cells is associated with reduced WNV pathogenesis. Our transcriptomic analysis revealed that IL-34 treatment upregulated markers characteristic of the DAM phenotype, including *TREM2*, *APOE*, *FABP5*, and *FTH1*, in HMC3 cells. Among these genes, the upregulation of *TREM2* was further confirmed via RT-qPCR, supporting the induction of a DAM-like transcriptional profile by IL-34.

These findings are consistent with the DAM signature, in which *TREM2* and *APOE* are representative upregulated genes (12). TREM2 is also a key regulator of the DAM transition (12, 23). Moreover, recent studies have suggested that TREM2 signaling is involved in microglial responses to IL-34 (21, 22). Thus, the IL-34–induced increase in *TREM2* expression observed in this study may represent one of the pathways through which IL-34 promotes a DAM-like cell state. IL-34–treated HMC3 cells suppressed WNV replication in co-culture with WNV-infected SH-SY5Y cells, but not under monoculture or supernatant transfer conditions, indicating that direct cell–cell contact is required for this antiviral effect. Microglia are essential for protection against WNV encephalitis, as their depletion results in increased viral titers and mortality in both *in vivo* and *ex vivo* models (26–28). Consistent with the known role of microglial phagocytosis in antiviral immunity (29, 30), the contact-dependent suppression observed in our study may involve enhanced recognition or clearance of WNV-infected neurons, rather than paracrine signaling. In addition, the close spatial association of TMEM119-positive microglia and SPP1-positive cells with WNV-infected cells, as observed via immunofluorescence, further supports this possibility. However, the precise molecular mechanisms underlying this contact-dependent suppression of viral replication remain to be elucidated.

Although IL-34 and CSF-1 share CSF1R, only IL-34 showed protective effects in WNV-infected mice. This difference suggests that general activation or expansion of TMEM119-positive microglia through CSF1R is not sufficient to suppress the progression of WNV disease. Indeed, CSF-1 increased TMEM119 immunoreactivity but failed to induce CD11c- or SPP1-positive DAM-like cells, whereas IL-34 promoted DAM-like cell responses in both uninfected and WNV-infected mouse brains. IL-34 and CSF-1 can exert distinct effects on macrophage or microglial activation states despite sharing CSF1R (18, 31). Therefore, the protective effect of IL-34 may be attributable not simply to CSF1R-mediated microglial expansion, but rather to its ability to induce a DAM-like state characterized by CD11c and SPP1 expression. However, the mechanisms by which IL-34, but not CSF-1, promotes this DAM-like response during WNV infection remain to be clarified.

This study had certain limitations. Most notably, the molecular mechanisms by which IL-34–treated microglia suppress WNV replication through direct cell–cell contact remain to be identified. In addition, we did not examine the transcriptomic profile of microglia from IL-34–treated WNV-infected mice. Therefore, it remains unclear whether the DAM-associated transcriptional signature observed *in vitro* and in uninfected mice is also induced during WNV infection. Furthermore, the transcriptomic analysis was performed using the HMC3 cell line rather than primary microglia, which may not fully recapitulate the *in vivo* phenotype of DAM. Finally, whether inhibition of CSF1R or IL-34 signaling affects DAM induction and viral replication during WNV infection remains to be investigated.

In conclusion, this study demonstrates that IL-34 promotes the activation of DAM-like cells and enhances protective responses in WNV infection both *in vitro* and *in vivo*. The induction of DAM-like cells by IL-34 provides novel insights into potential therapeutic strategies aimed at modulating microglial responses in WNV encephalitis. Future studies should focus on elucidating the molecular mechanisms underlying IL-34–mediated DAM induction, including the possible involvement of CSF1R and TREM2 signaling, and evaluate its therapeutic potential in other neurotropic viral infections.

## MATERIALS AND METHODS

### Ethics statement

All animal experiments were performed following the basic guidelines for animal experiments prescribed by the MEXT, Japan. The President of Hokkaido University approved all animal experiments after review by the Institutional Animal Care and Use Committee of Hokkaido University (approval no. 23-0128).

### Cells and culture conditions

Human neuroblastoma cell line (SH-SY5Y) was obtained from the European Collection of Authenticated Cell Cultures (ECACC; catalog no. 94030304) and cultured in Dulbecco’s Modified Eagle Medium/Nutrient Mixture F-12 (DMEM/F-12; Wako, Osaka, Japan). Human microglial clone 3 (HMC3) was purchased from American Type Culture Collection (Manassas, VA, USA; catalog no. CRL-3304). African green monkey kidney epithelial cells (Vero) were obtained from the Japanese Collection of Research Bioresources Cell Bank (Osaka, Japan; catalog no. JCRB0111). HMC3 and Vero cells were grown in Eagle’s Minimum Essential Medium (MEM; Wako). All culture media were supplemented with 10% fetal bovine serum (FBS), 100 IU/mL penicillin, and 100 μg/mL streptomycin. All cells were maintained in a humidified incubator at 37 °C with 5% CO2 and passaged at 80%–90% confluency.

### Recombinant proteins and treatment

Recombinant human IL-34 (catalog no. 577902), recombinant mouse IL-34 (catalog no. 577602), and recombinant mouse CSF-1 (M-CSF; catalog no. 576402) were purchased from BioLegend (San Diego, CA, USA). HMC3 cells were treated with recombinant human IL-34 at a final concentration of 200 ng/mL for 24 h prior to further cell culture experiments. For animal experiments, recombinant mouse IL-34 or CSF-1 (100 ng) prepared in PBS (10 μL) was administered via intracranial injection under isoflurane anesthesia one day after viral inoculation. Vehicle-treated control mice received 10 μL of PBS via the same route.

### Virus preparation for infection

WNV NY99 6-LP strain was propagated in Vero cells (7) and stored at −80 °C until use. To assess the susceptibility of HMC3 and SH-SY5Y cells to WNV, the cells were infected at a multiplicity of infection (MOI) of 0.01 or 0.1 and analyzed at 24 or 48 hpi. Subsequent cell infection experiments were performed at an MOI of 0.1. For animal experiments, an inoculation dose of 10 pfu per mouse was used, as described below.

For co-culture experiments, HMC3 cells were pre-treated with recombinant human IL-34 prior to co-culture. IL-34–treated HMC3 cells were co-cultured with WNV-infected SH-SY5Y cells at a ratio of 1:1. For supernatant transfer experiments, conditioned medium collected from IL-34–treated HMC3 cells was diluted 1:1 with DMEM/Ham’s F-12 medium and transferred to WNV-infected SH-SY5Y cell culture. Viral titers were measured at 12 or 24 hpi via plaque assay. All experiments using live WNV were performed under Biosafety Level 3 conditions in accordance with institutional guidelines.

### Animal experiments

Five-week-old female C57BL/6JmsSlc mice were purchased from Japan SLC Inc. (Hamamatsu, Japan) and housed under standard conditions. WNV inoculation was performed via intracranial injection of 10 pfu per mouse in Hank’s Balanced Salt Solution (HBSS) under isoflurane anesthesia. The virus was administered into the left frontal lobe of the brain. Mice were monitored daily and body weights were recorded throughout the experiment. The brain tissue was collected at 7 dpi for further analysis. Humane endpoints were defined as a 20% loss in body weight or an inability to walk or reach food and water because of disease onset.

### Plaque assay

The brain tissue was individually weighed, manually homogenized, and prepared as 10% (w/v) suspension in PBS. The suspension was centrifuged at 3,000 × *g* for 5 min at 4 °C, and supernatant was collected and stored at −80 °C until use. Vero cells were seeded in 24-well plates and infected the following day with 10-fold serial dilutions of brain homogenate or cell culture supernatant. After 1 h of adsorption with shaking at 37 °C, cells were overlaid with MEM containing 5% FBS and 1.25% methylcellulose and incubated for 4 days at 37 °C with 5% CO2. Plaques were visualized by removing the overlay medium and staining with 0.25% crystal violet in 10% formalin. Viral titers were calculated and expressed as log10 pfu/mL.

### Immunofluorescence staining and immunohistochemistry

For staining of the brain tissue, mice were euthanized via isoflurane over-inhalation, transcardially perfused with PBS and 4% paraformaldehyde (PFA), and their brain was post-fixed in formalin for 48 h followed by paraffin embedding. Coronal sections (4 μm) were deparaffinized, rehydrated, and subjected to antigen retrieval using 10 mM citrate buffer (pH 6.0). The sections were blocked with 10% goat serum and incubated with primary antibodies in Can Get Signal solution A (Toyobo, Osaka, Japan) overnight at 4 °C. For cell staining, cells were fixed with 4% PFA for 10 min, blocked with 1% BSA-PBS, and incubated with primary antibodies overnight at 4 °C.

The following primary antibodies were used: anti-WNV E-protein mouse monoclonal antibody (1:100 for tissue, 1:200 for cells; Merck Millipore, catalog no. MAB8151), anti-TMEM119 rabbit monoclonal antibody (1:1000; Abcam, Cambridge, UK, catalog no. ab209064) for tissue or rabbit polyclonal antibody (1:200; Proteintech, Rosemont, IL, USA, catalog no. 27585-1-AP) for cells, anti-CD11c rabbit polyclonal antibody (1:100; Proteintech, catalog no. 17342-1-AP), anti-cleaved caspase-3 rabbit polyclonal antibody (1:200; Proteintech, catalog no. 25128-1-AP), and anti-SPP1 rabbit monoclonal antibody (E9Z1D, 1:200; Cell Signaling Technology, catalog no. 88742). For immunohistochemistry, the sections were incubated with secondary antibody for 1 h at room temperature, visualized using a substrate kit (Nichirei, Tokyo, Japan), and counterstained with hematoxylin. Images were captured using a BX60 microscope with cellSens software (Olympus, Tokyo, Japan). For immunofluorescence, Alexa Fluor 488- and 555-conjugated secondary antibodies (1:1,000; Thermo Fisher Scientific, Waltham, MA, USA; catalog no. A-11001 and A-21428) and Hoechst 33342 (1:1,000; WAKO; catalog no. 346-07951) were applied for 1 h at room temperature. Anti-TMEM119 antibody for triple immunofluorescence staining was directly labeled using Zenon rabbit IgG labeling kits (Thermo Fisher Scientific; catalog no. Z25608). Images of the brain tissue and cells were acquired using LSM 700 and LSM 800 confocal microscopes, respectively, with ZEN software (Carl Zeiss, Oberkochen, Germany). Cells positive for the stain were manually counted in at least three biological replicates.

### Flow cytometry

Mouse brains were dissociated into single-cell suspensions as previously described (15). Briefly, fresh whole brains were collected in cold HBSS and mechanically dissociated in HBSS containing DNase I (Roche). Cell suspensions were filtered through a 70-micrometer cell strainer (Greiner Bio-One, Milan, Italy), and myelin was removed using 30% Percoll solution (GE Healthcare, Uppsala, Sweden). Single-cell suspensions were blocked with 1% BSA in PBS for at least 10 min and incubated with APC-conjugated anti-CD11c mouse monoclonal antibody (0.25 μg/test; Thermo Fisher Scientific; catalog no. 17-0114-82) and Alexa Fluor 488-conjugated anti-TMEM119 rabbit monoclonal antibody (1/500; Abcam; catalog no. ab225497) for 30 min. Fixable Viability Dye eFluor 780 (1:1000; Thermo Fisher Scientific; catalog no. 65-0865-14) was included for the exclusion of dead cells. The cells were fixed with 4% PFA for 15 min and filtered through a 40-micrometer cell strainer (Greiner Bio-One). Data were acquired using a BD FACSLyric flow cytometer and analyzed with BD FACSuite software v.1.4 (BD Biosciences). The gating strategy discriminated doublets based on forward scatter area and height, separated debris and erythrocytes based on forward scatter and side scatter area, and excluded dead cells prior to analysis of CD11c and TMEM119 expression.

### Bulk RNA sequencing

Total RNA was extracted from HMC3 cells using IsogenII (Nippon Gene, Tokyo, Japan) with the Direct-zol RNA Miniprep Kit (Zymo Research, Irvine, CA, USA) according to the manufacturer’s instructions. HMC3 cells were treated with IL-34 or vehicle for 24 h in biological triplicates. RNA quality and quantity were assessed prior to library preparation. RNA sequencing libraries were prepared and sequenced on an Illumina NovaSeq X Plus platform (Illumina, San Diego, CA, USA) with 150 bp paired-end reads at a depth of approximately 40 million reads per sample (20 million pairs). All procedures were performed at Rhelixa (Tokyo, Japan).

Raw sequencing reads were processed and aligned to the human reference genome (GRCh38) using the RaNAseq web-based pipeline (https://ranaseq.eu). Raw count matrices were downloaded and imported into R version 4.5.3 for downstream analysis. Principal component analysis (PCA) was performed to assess sample quality. One IL-34–treated sample was identified as an outlier and excluded from downstream analysis, resulting in *n* = 2 for the IL-34–treated group and *n* = 3 for the vehicle-treated group. Differential gene expression analysis was performed using the DESeq2 package (v.1.50.2) with a minimum read-count threshold of 10. DEGs were identified using thresholds set at log2 fold change >0.25 and adjusted *P*-value <0.05. GO biological process enrichment analysis was performed using the clusterProfiler package (v.4.18.4) with the org.Hs.eg.db annotation database (v.3.22.0). Redundant GO terms were removed using the simplify() function with a similarity cutoff of 0.5. Heatmaps and volcano plots were generated using the pheatmap (v.1.0.13) and ggplot2 (v.4.0.2) packages, respectively. Gene labels in volcano plots were annotated using the ggrepel package (v.0.9.8). GO enrichment results were visualized using the enrichplot package (v.1.30.5). Data manipulation was performed using the dplyr package (v.1.2.1).

### RNA isolation and RT-qPCR

For HMC3 cells, total RNA was extracted as described above. *CD11c* expression was quantified using the Thunderbird Probe One-Step qRT-PCR Kit (Toyobo, Osaka, Japan) with a predesigned probe-based assay targeting *ITGAX* (*CD11c*; Hs.PT.58.40875739; Integrated DNA Technologies, Coralville, IA, USA), with *ACTB* (Hs.PT.39a.22214847; Integrated DNA Technologies) used as the reference gene. To quantify *TREM2* and *SPP1* expression via SYBR Green-based assay, complementary DNA (cDNA) was synthesized from total RNA using the PrimeScript II First-Strand cDNA Synthesis Kit (Takara Bio, Shiga, Japan) (30 °C for 10 min, 42 °C for 60 min, and 95 °C for 5 min), and quantified using the KAPA SYBR FAST qPCR Master Mix (2X) (Kapa Biosystems, Wilmington, MA, USA) on a 7500 Fast Real-Time PCR System (Thermo Fisher Scientific), with the following cycling conditions: 95 °C for 3 min, followed by 40 cycles of 95 °C for 3 s and 60 °C for 25 s. *GAPDH* served as the reference gene. In mouse brain tissue, TMEM119-sorted microglia were lysed, and total RNA was extracted using the Direct-zol RNA MiniPrep Kit (Zymo Research) according to the manufacturer’s protocol. cDNA synthesis and qPCR were performed as described above for SYBR Green-based assays, with *CD11c* expression quantified using *GAPDH* as the reference gene. For all assays, relative gene expression was calculated using the delta-delta cycle threshold (ΔΔCt) method. Primer sequences are listed in Table S1.

### Statistical analysis

All statistical analyses were performed using GraphPad Prism version 10. Data are presented as the mean ± standard deviation (SD). For experiments comparing two independent groups, an unpaired two-tailed Student’s *t*-test was used. For comparisons among multiple groups, one-way analysis of variance (ANOVA) with Tukey’s or Dunnett’s multiple comparisons post-hoc test was applied, as appropriate. Survival curves were estimated using the Kaplan–Meier method, and differences between groups were assessed using the log-rank test. *P*-values <0.05 were considered to indicate statistically significant difference.

### Data availability

Bulk RNA-seq data are available in the DNA Data Bank of Japan (DDBJ) under BioProject accession number PRJDB42796. All other data supporting the findings of this study are available from the corresponding author upon reasonable request.

## ACKNOWLEDGMENTS

This study was supported in part by the Japan Society for the Promotion of Science (JSPS) KAKENHI Grant Numbers 24K01917, 24KJ0264, 25K22699, 25K09430, and 26KJ0446; the Japan Agency for Medical Research and Development (AMED) under Grants JP25wm0325077, JP223fa627005, JP20wm0125008, and 25wm0325073 ; the Naito Foundation; the Shionogi infectious disease research promotion foundation; the Takeda Science Foundation; a research grant from the Astellas Foundation for Research on Metabolic Disorders; JST SPRING (JPMJSP2119); JST Moonshot R&D (JPMJMS2025); and the World-leading Innovative and Smart Education (WISE) program (1801) from the Ministry of Education, Culture, Sports, and Technology, Japan

Passawat Thamhamakin: Conceptualization, Methodology, Investigation, Formal analysis, Writing – original draft, Writing – review & editing. Keisuke Maezono: Methodology, Investigation, Funding acquisition. Haruto Eguchi: Methodology, Investigation, Funding acquisition. Thi Ngoc Thuy Duong and Savit Promwattanapan: Methodology, Investigation. Michihito Sasaki: Resourses, Writing – review & editing. Hiroaki Kariwa: Resources, Supervision. Orba Yasuko: Resources, Writing – review & editing, Funding acquisition. Shintaro Kobayashi: Conceptualization, Resources, Funding acquisition, Supervision, Writing – review & editing. All authors read and approved the final manuscript.

---

**FIG S1** Additional transcriptomic analysis of interleukin (IL-34)–treated HMC3 cells. (A) Dot plot of gene ontology (GO) biological process enrichment analysis of upregulated genes in IL-34–treated HMC3 cells. Dot size represents gene count, and color represents adjusted *P*-value. (B) Dot plot of GO biological process enrichment analysis of downregulated genes in IL-34–treated HMC3 cells. Dot size represents gene count, and color represents adjusted *P*-value. (C) Heatmap of the top 10 differentially expressed genes (DEGs) ranked by adjusted *P*-value in IL-34–treated to vehicle-treated HMC3 cells. Color scale represents row-normalized expression values.

**FIG S2** Susceptibility of HMC3 and SH-SY5Y cells to West Nile virus (WNV) infection. Representative immunofluorescence images of WNV antigen in HMC3 and SH-SY5Y cells infected with WNV at a multiplicity of infection (MOI) of 0.01 or 0.1, or mock-infected, at 24 and 48 post-infection (hpi). Scale bar, 300 μm.

**FIG S3** Effect of colony-stimulating factor 1 (CSF-1) administration on microglia in uninfected mice. (A) Representative immunohistochemistry images of TMEM119 in brain sections from mice treated with vehicle or CSF-1. Scale bar, 50 μm. (B) Quantification of the average area of TMEM119-positive cells representing microglial size. Data represent the mean ± SD from three fields per mouse (*n* = 3 mice per group). (C) Quantification of CD11c-positive cells as a percentage of total live cells. Data represent the mean ± SD (*n* = 3 mice per group). (D) Representative immunofluorescence images of brain sections from vehicle- and CSF-1–treated mice stained for SPP1. Scale bar, 100 μm. Statistical significance was determined via Student’s *t*-test. ns, not significant; ****, *P* < 0.0001.

**FIG S4** Effect of colony-stimulating factor 1 (CSF-1) treatment on body weight, viral titer, and neuronal apoptosis in West Nile virus (WNV)-infected mice. (A) Percentage change in body weight at 7 days post-infection (dpi) relative to initial body weight in WNV-infected mice treated with vehicle or CSF-1 (*n* ≥ 19 per group). (B) WNV titers in brain tissues collected at 7 dpi from WNV-infected mice treated with vehicle or CSF-1, determined via plaque assay on Vero cells. Data are shown as log10 (pfu/mL) (*n* = 8 per group). The dotted line indicates the limit of detection. (B) Representative immunofluorescence images of cleaved caspase-3 (red) in brain sections from WNV-infected mice treated with vehicle (WNV+Vehicle) or CSF-1 (WNV+CSF-1). Scale bar, 100 μm. (D) Quantification of cleaved caspase-3-positive cells per counting area. Data are presented as the mean ± SD from five fields per mouse (*n* = 5 per group). Statistical significance was determined via Student’s *t*-test. ns, not significant.

**FIG S5** Examination of the effects of colony-stimulating factor 1 (CSF-1) on CD11c-positive microglia in West Nile virus (WNV)-infected mouse brains. (A) Representative immunofluorescence images of WNV antigen (green) and CD11c (red) in brain sections from WNV-infected mice treated with vehicle (WNV+Vehicle) or CSF-1 (WNV+CSF-1). White arrowheads indicate CD11c-positive cells in contact with WNV-infected cells. Scale bar, 50 μm. (B) Quantification of CD11c-positive cells per counting area from five fields per mouse (*n* = 4 per group). (C) Quantification of the percentage of CD11c-positive cells in contact with WNV-infected cells out of total CD11c-positive cells from five fields per mouse (*n* = 4 per group). Data are presented as the mean ± SD. Statistical significance was determined via Student’s *t*-test. ns, not significant.

## REFERENCES

1. Samuel MA, Diamond MS. 2006. Pathogenesis of West Nile Virus Infection: a Balance between Virulence, Innate and Adaptive Immunity, and Viral Evasion. J Virol 80:9349–9360.

2. Lim SM, Koraka P, Osterhaus ADME, Martina BEE. 2011. West Nile Virus: Immunity and Pathogenesis. Viruses 3:811–828.

3. Colpitts TM, Conway MJ, Montgomery RR, Fikrig E. 2012. West Nile Virus: Biology, Transmission, and Human Infection. Clin Microbiol Rev 25:635–648.

4. Sejvar JJ, Haddad MB, Tierney BC, Campbell GL, Marfin AA, Van Gerpen JA, Fleischauer A, Leis AA, Stokic DS, Petersen LR. 2003. Neurologic manifestations and outcome of West Nile virus infection. Jama 290:511–515.

5. McDonald E, Mathis S, Martin SW, Erin Staples J, Fischer M, Lindsey NP. 2021. Surveillance for West Nile virus disease — United States, 2009–2018. Am J Transplant 21:1959–1974.

6. Clarke P, Leser JS, Quick ED, Dionne KR, Beckham JD, Tyler KL. 2014. Death Receptor-Mediated Apoptotic Signaling Is Activated in the Brain following Infection with West Nile Virus in the Absence of a Peripheral Immune Response. J Virol 88:1080–1089.

7. Kobayashi S, Orba Y, Yamaguchi H, Kimura T, Sawa H. 2012. Accumulation of ubiquitinated proteins is related to West Nile virus-induced neuronal apoptosis. Neuropathology 32:398–405.

8. Stonedahl S, Clarke P, Tyler KL. 2020. The role of microglia during West Nile virus infection of the central nervous system. Vaccines 8:485.

9. Ulbert S. 2019. West Nile virus vaccines – current situation and future directions. Hum Vaccines Immunother 15:2337–2342.

10. Gould CV, Staples JE, Huang CY-H, Brault AC, Nett RJ. 2023. Combating West Nile Virus Disease — Time to Revisit Vaccination. N Engl J Med 388:1633–1636.

11. Waltl I, Kalinke U. 2022. Beneficial and detrimental functions of microglia during viral encephalitis. Trends Neurosci 45:158–170.

12. Keren-Shaul H, Spinrad A, Weiner A, Matcovitch-Natan O, Dvir-Szternfeld R, Ulland TK, David E, Baruch K, Lara-Astaiso D, Toth B. 2017. A unique microglia type associated with restricting development of Alzheimer’s disease. Cell 169:1276–1290.

13. Deczkowska A, Keren-Shaul H, Weiner A, Colonna M, Schwartz M, Amit I. 2018. Disease-associated microglia: a universal immune sensor of neurodegeneration. Cell 173:1073–1081.

14. Cheng Y-H, Ho MS. 2025. Disease-associated microglia in neurodegenerative diseases: Friend or foe? PLoS Biol 23:e3003426.

15. Thammahakin P, Maezono K, Maekawa N, Kariwa H, Kobayashi S. 2023. Detection of disease-associated microglia among various microglia phenotypes induced by West Nile virus infection in mice. J Neurovirol 29:367–375.

16. Elmore MR, Najafi AR, Koike MA, Dagher NN, Spangenberg EE, Rice RA, Kitazawa M, Matusow B, Nguyen H, West BL. 2014. Colony-stimulating factor 1 receptor signaling is necessary for microglia viability, unmasking a microglia progenitor cell in the adult brain. Neuron 82:380–397.

17. Wang Y, Szretter KJ, Vermi W, Gilfillan S, Rossini C, Cella M, Barrow AD, Diamond MS, Colonna M. 2012. IL-34 is a tissue-restricted ligand of CSF1R required for the development of Langerhans cells and microglia. Nat Immunol 13:753–760.

18. Easley-Neal C, Foreman O, Sharma N, Zarrin AA, Weimer RM. 2019. CSF1R ligands IL-34 and CSF1 are differentially required for microglia development and maintenance in white and gray matter brain regions. Front Immunol 10:2199.

19. Devlin BA, Nguyen DM, Ribeiro D, Grullon G, Clark MJ, Finn A, Ceasrine AM, Oxendine S, Deja M, Shah A. 2025. Excitatory-neuron-derived interleukin-34 supports cortical developmental microglia function. Immunity 58:1948–1965.

20. Mizuno T, Doi Y, Mizoguchi H, Jin S, Noda M, Sonobe Y, Takeuchi H, Suzumura A. 2011. Interleukin-34 selectively enhances the neuroprotective effects of microglia to attenuate oligomeric amyloid-β neurotoxicity. Am J Pathol 179:2016–2027.

21. Shang J, Xu Y, Pu S, Sun X, Gao X. 2023. Role of IL-34 and its receptors in inflammatory diseases. Cytokine 171:156348.

22. Xie X, Zhang W, Xiao M, Wei T, Qiu Y, Qiu J, Wang H, Qiu Z, Zhang S, Pan Y. 2023. TREM2 acts as a receptor for IL-34 to suppress acute myeloid leukemia in mice. Blood J Am Soc Hematol 141:3184–3198.

23. Krasemann S, Madore C, Cialic R, Baufeld C, Calcagno N, El Fatimy R, Beckers L, O’loughlin E, Xu Y, Fanek Z. 2017. The TREM2-APOE pathway drives the transcriptional phenotype of dysfunctional microglia in neurodegenerative diseases. Immunity 47:566–581.

24. Berglund R, Cheng Y, Piket E, Adzemovic MZ, Zeitelhofer M, Olsson T, Guerreiro-Cacais AO, Jagodic M. 2024. The aging mouse CNS is protected by an autophagy-dependent microglia population promoted by IL-34. Nat Commun 15:383.

25. Samuel MA, Morrey JD, Diamond MS. 2007. Caspase 3-Dependent Cell Death of Neurons Contributes to the Pathogenesis of West Nile Virus Encephalitis. J Virol 81:2614–2623.

26. Funk KE, Klein RS. 2019. CSF1R antagonism limits local restimulation of antiviral CD8+ T cells during viral encephalitis. J Neuroinflammation 16:22.

27. Stonedahl S, Leser JS, Clarke P, Tyler KL. 2022. Depletion of Microglia in an *Ex Vivo* Brain Slice Culture Model of West Nile Virus Infection Leads to Increased Viral Titers and Cell Death. Microbiol Spectr 10:e00685–22.

28. Chhatbar C, Prinz M. 2021. The roles of microglia in viral encephalitis: from sensome to therapeutic targeting. Cell Mol Immunol 18:250–258.

29. Quick ED, Leser JS, Clarke P, Tyler KL. 2014. Activation of Intrinsic Immune Responses and Microglial Phagocytosis in an *Ex Vivo* Spinal Cord Slice Culture Model of West Nile Virus Infection. J Virol 88:13005–13014.

30. Hatton CF, Duncan CJ. 2019. Microglia are essential to protective antiviral immunity: lessons from mouse models of viral encephalitis. Front Immunol 10:2656.

31. Boulakirba S, Pfeifer A, Mhaidly R, Obba S, Goulard M, Schmitt T, Chaintreuil P, Calleja A, Furstoss N, Orange F. 2018. IL-34 and CSF-1 display an equivalent macrophage differentiation ability but a different polarization potential. Sci Rep 8:256.

